# Tspan15 depletion in mice affects social investigation and aggressive behavior

**DOI:** 10.1101/2024.05.16.594514

**Authors:** Daniele Stajano, Sandra Freitag, Paul Saftig, Kira V. Gromova, Matthias Kneussel

## Abstract

Tetraspanins are transmembrane proteins that form tetraspanin-enriched microdomains at biological membranes and contribute to the formation of functional protein complexes involved in various cellular processes. Tspan7 and Tspan15 are highly expressed in the hippocampus, a brain region that contributes to the regulation of social aggression. While Tspan7 has been observed to modulate cognition and behavior in rodents, the function of Tspan15 in this context is still unknown. In this study, we tested preference for social novelty, territorial and aggressive behavior in mice lacking Tspan15 gene expression. We report that male Tspan15-KO mice exhibit an increased motivation for social investigation and show increased aggression compared to WT littermates, regardless of familiarity with the partner mouse. Our data suggest a possible role of Tspan15 as a molecular modulator of certain neuronal circuits that control aggressive behavior.

## Introduction

Tetraspanins (Tspans) represent an evolutionarily conserved family of small (200-300 amino acids) transmembrane proteins mainly found at intracellular vesicle membranes, at the plasma membrane (Charrin *et al*, 2009) or on extracellular vesicles (Andreu & Yanez-Mo, 2014). On biological membranes, tetraspanins assemble with each other and organize their interaction partners into functional complexes on platforms called tetraspanin-enriched microdomains (TEMs) (Murru *et al*, 2018; van Deventer *et al*, 2017). Within the TEMs, different interaction partners of tetraspanins can be stabilized in close proximity to each other, which is why tetraspanins are also referred to as molecular mediators (van Deventer *et al*., 2017). However, individual tetraspanins show very different expression patterns in various cell types including neurons of the brain (Murru *et al*., 2018). Tspan7 and Tspan15 are strongly expressed in the hippocampus of the central nervous system (Bassani *et al*, 2012; Prox *et al*, 2012), which is associated with the disinhibition of motivated behaviors such as social aggression (Leroy *et al*, 2018). Tspan15 has been extensively studied as an interaction partner of A disintegrin and metalloprotease 10 (ADAM10), promoting its trafficking to the plasma membrane and its activity through the formation of a molecular scissor complex (Seipold *et al*, 2018). In addition, Tspan15 was recently reported to modulate intercellular communication by facilitating contacts between neuron-derived extracellular vesicles and target neurons (Stajano *et al*, 2023). However, while Tspan7 has been associated with behavioral phenotypes reminiscent of autism and intellectual disability, including impaired sociability (Bassani *et al*., 2012; Bassani & Passafaro, 2012; Becic *et al*, 2021; Murru *et al*, 2021), little is known about the role of Tspan15 in this context.

For several animal species, social contact is essential for survival, and aggressive behavior represents a motivated social behavior that occurs when physical interactions take place to overcome critical situations, such as establishing social hierarchy, competing for food, and defending offspring and territory (Takahashi & Miczek, 2014). Whether or not to engage in a fight depends on a cost-benefit analysis, and if this calculation is impaired or distorted, it can lead to escalated aggressive behavior (Takahashi & Miczek, 2014). Several human neurological and psychiatric disorders are characterized by exaggerated aggressiveness, such as Fragile X syndrome (Lozano *et al*, 2014), but the molecular players involved in these biological processes are not yet fully understood.

Here, we report a novel function of Tspan15 in the modulation of social investigation and aggressive behavior in mice.

## Results and Discussion

### Depletion of Tspan15 leads to alterations in social investigation and aggressive behavior

Members of the tetraspanin family, such as Tspan7, have been reported to regulate cognition and sociability in rodents (Murru *et al*., 2021). In the present study, we first aimed to investigate behavioral consequences of Tspan15 depletion using Tspan15 knockout mice in comparison to wild-type littermates. For this purpose, male littermates were housed together as single pairs and their behavior was examined using different paradigms. In an initial open field test, Tspan15 knockout animals showed normal locomotor and exploratory activity compared to wild-type controls (not shown). Likewise, in an elevated-plus maze test, Tspan-15 wild-type and knockout littermates also showed the same behavior with no difference in anxiety levels (not shown).

We also tested the preference for social novelty comparing the social investigation of a test mouse with familiar and unfamiliar animals (Figure 1A). In this series of experiments, wild-type and Tspan15-KO mice did not show an altered preference index towards a familiar or an unfamiliar mouse (Figure 1B). However, the duration of time spent in close proximity to other mice (social time) was increased in knockout animals lacking Tspan15 gene expression (Figure 1C), suggesting that Tspan15 KOs may have a higher motivation during social investigation.

**Figure 1.**
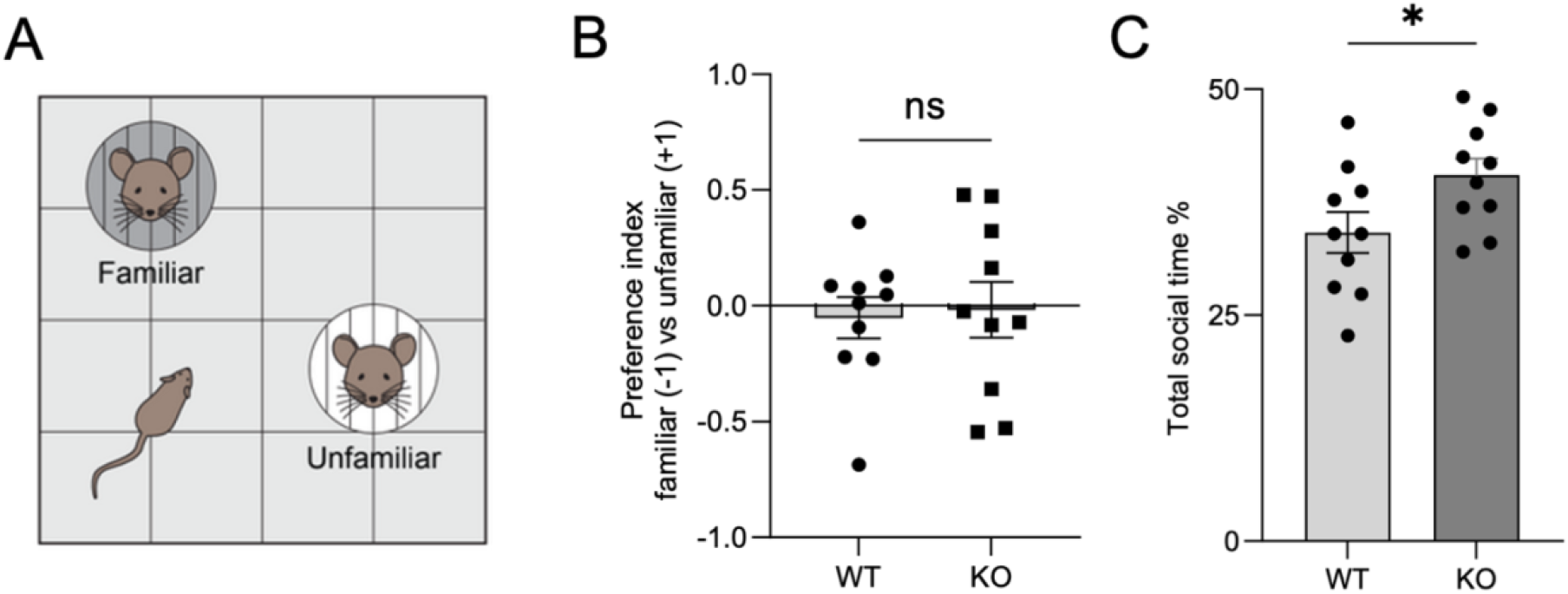
Depletion of Tspan15 expression increases social interest in male mice. (A) Schematics of social preference test. (B) Quantification of social preference index. Wild-type (WT), Tspan15 knockout (KO). Student’s t-test. ns=not significant. (C) Quantification of percentage of social time, expressing the total time in proximity to any partner mouse. Student`s t-test, p: 0.0431. n=10 male mice per genotype from 4 cohorts. Data are expressed as individual data points ± SEM.

In addition, a urine marking test was carried out to assess possible changes in territorial behavior (Drickamer, 2001) (Figure 2A). Wild-type and Tspan15 knockout mice left a similar number of urine traces throughout the arena (Figure 2B). Similarly, there was no significant difference in the number of urine traces in a quadrant of the arena where three female mice were housed (Figure 2C). Thus, we conclude that depletion of Tspan15 does not affect the territorial behavior of males.

**Figure 2.**
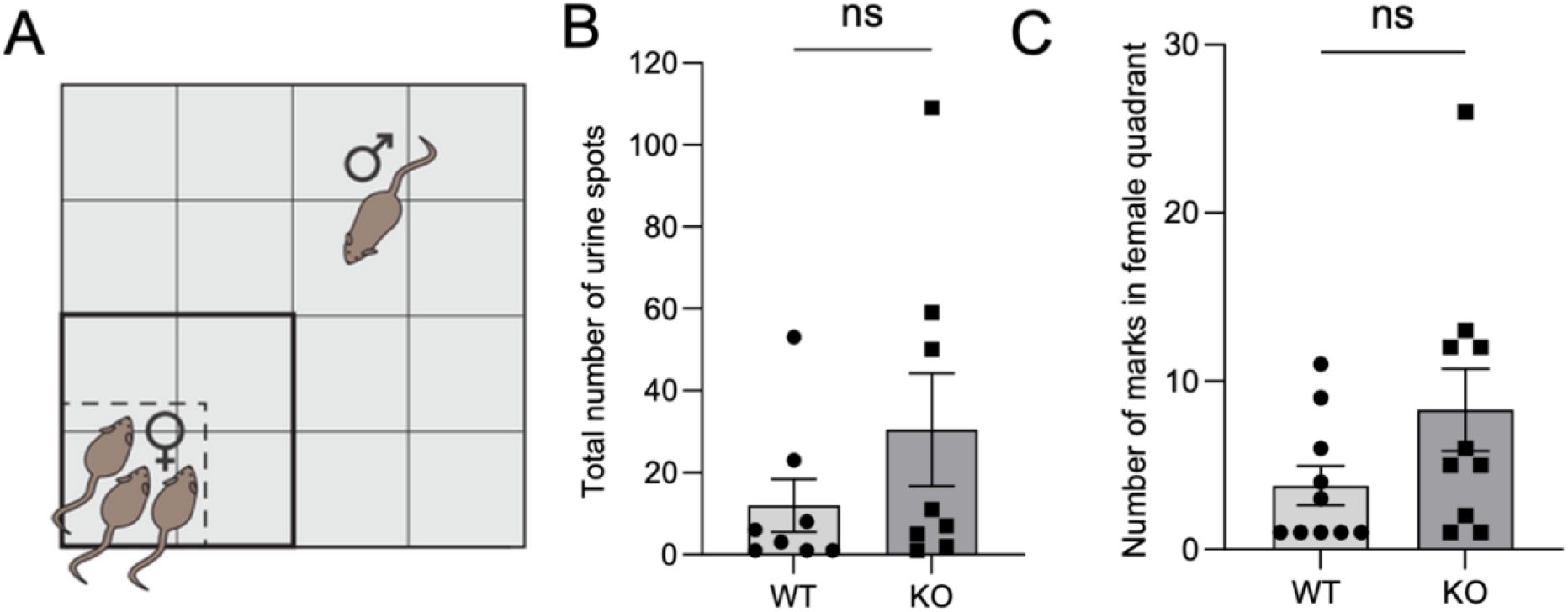
Wild-type and Tspan15 knockout mice show similar territorial behavior. (A) Schematic of urine marking test. ♀: female partner mice. ♂: focal wild-type (WT) or Tspan15 knockout (KO) male mice. (B) Quantification of total urine marks in the arena. Mann Whitney U test. ns= not significant. (C) Quantification of urine marks released in the female-hosting quadrant. Mann Whitney U test. ns= not significant. n=10 male mice per genotype from 4 cohorts. Data are expressed as individual data points ± SEM.

We further assessed aggressive behavior between cage mates in the home cage environment with immediate sampling after the cage change (Figure 3A). Since the animals had interacted before and during the experiment (dependent samples), a Wilcoxon test was used to assess statistical significance. Under these conditions, we observed that Tspan15-KO mice showed a higher percentage of aggression than their wild-type cage mates (Figure 3B).

**Figure 3.**
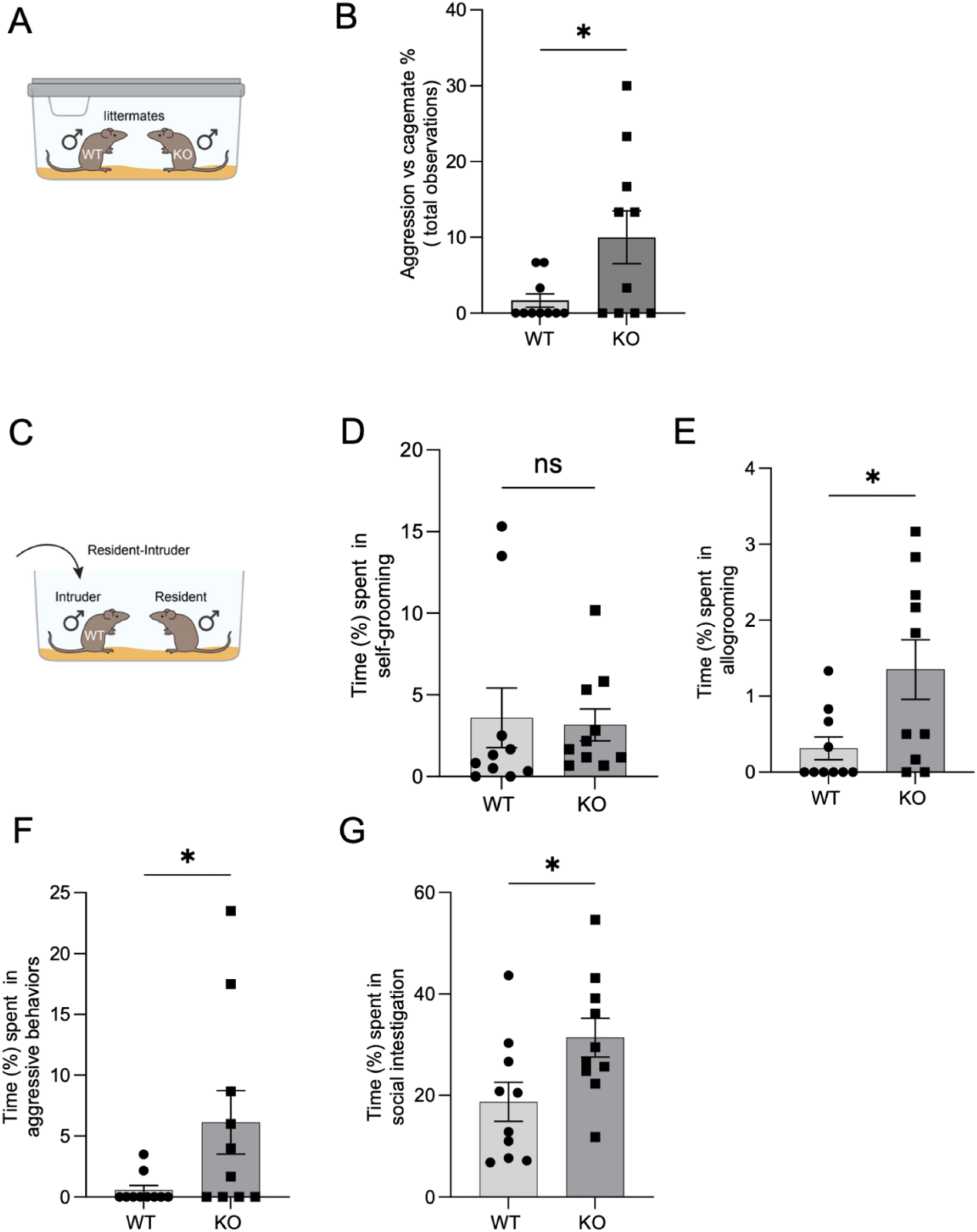
The loss of Tspan15 expression leads to enhanced aggression. (A) Schematic of experimental design. (B) Quantification of the percentage of aggression towards the cage brother in the home cage when the bedding was changed. Wilcoxon matched-pairs signed rank test, p: 0.0469. (C) Schematic representation of the test setup for the resident intruder test. (D-G) Quantifications from Resident-Intruder test. (D) Duration of self-grooming. Mann-Whitney U test. ns= not significant. (E) Duration of allogrooming. Mann-Whitney U test, p: 0.0423. (F) Duration of aggressive episodes versus the intruder. Mann-Whitney U test. p: 0.0412. (G) Duration of social investigation. Mann-Whitney U test. p: 0.0373. n=10 male mice per genotype from 4 cohorts. Data are expressed as individual data points ± SEM.========

To determine whether this observation might manifest also in an independent experiment with unfamiliar mice, we aimed to conduct a resident intruder test, a standardized method for measuring offensive aggression and defensive behavior in a semi-natural environment (Figure 3C). Interestingly, although the wild-type and Tspan15 KO animals did not differ in the percentage of time spent on self-grooming (Figure 3D), the KO animals spent significantly more time on allogrooming, which could be related to the observed higher motivation during the social investigation (compare with Figure 1C). Notably, Tspan15 knockout animals spent significantly more time in aggressive behavior (Figure 3F) and in social investigation versus the WT intruder (Figure 3G).

Taken together, these data show that depletion of Tspan15 appears to lead to increased aggression and higher motivation in social investigations.

Tetraspanins regulate various membrane proteins in different cell types. We have previously shown that Tspan15 is a component of neuronally generated extracellular vesicles that are transmitted intercellularly between neurons (Stajano *et al*., 2023). Future studies at the molecular and cellular level will have to clarify how Tspan15 contributes to the described behavioral changes. In fact, several neurological and psychiatric disorders in patients are associated with exaggerated aggression, such as fragile X syndrome (Lozano *et al*., 2014). Whether Tspan15 is related to these diseases will be a question to be answered in the future.

## Materials and Methods

### Animals

Tspan15 KO mice, described previously (Seipold *et al*., 2018), were housed and bred at the Center of Molecular Neurobiology Hamburg animal facility. All procedures were fulfilled by the European Communities Council Directive (2010/63/EU) on the protection of experimental animals and guiding principles expressed by the German Animal Welfare Act. All protocols were carried out with the approval of the ethics committee of the City of Hamburg (Behörde für Gesundheit und Verbraucherschutz, Fachbereich Veterinärwesen; No. 100/2021).

### Genotyping

Genomic DNA was extracted from tail applying QuickExtract DNA Extraction Buffer (Biozym Scientific GmbH, Hessisch Oldendorf) and performing PCR with the following primers: Tspan15 exon2 sense: 5’-AAGCTTGGGCTATAGGCATACACC-3’; antisense 5’-GGATCCCCAGCATCTCTGTGACAGC-3’. The presence of wild-type and knockout alleles was shown by two different amplicons, respectively of 600bp and/or 500bp, on a 1% agarose gel.

### Statistics

All statistics were calculated using Prism (GraphPad, version 9.0.1). Graphs were generated using Prism and images were assembled with Adobe Illustrator 2021. Then data were tested for normality using Kolmogorov-Smirnov or Shapiro-Wilk tests. For normally distributed data, a two-tailed unpaired Student’s t-test was used according to the experimental design. For non-normally distributed data the Wilcoxon t-test or Mann-Whitney U test were used to test the statistical significance. Statistical significance was represented as follows: *p >0.05. ns: not significant Additional statistical information for single experiments is listed in the corresponding figure legends.

### Behavioral analysis

The Tspan15 KO line was backcrossed at least three generations and maintained in a C57BL/6 background. Focal animals were generated from heterozygous x heterozygous matings. Mice were housed in couples of same-sex littermates in a temperature (22 ± 1°C) and humidity (50 ± 5%)-controlled vivarium on an inverted 12h:12h light-dark cycle. Not more than two couples per litter were selected to avoid litter-effects. Animals had ad libitum access to food and water. Animals were 16-18 weeks old prior to behavioral testing, which was carried out during the dark phase of the cycle. Behavioral experiments were conducted on male mice. The number of mice used per genotype is provided in the corresponding figure legend for each experiment.

Mice used for behavioral tests were initially observed blind to genotype in the home cage as described (Freitag *et al*, 2003). In brief, observations were performed three times per day (morning, afternoon and late afternoon) in the home cage for 1 hour to assess the spontaneous home cage activity. Observations of spontaneous home activity were also performed following the cage change to evaluate the spontaneous social abilities between cagemates. The WT and Tspan15 KO cagemates were marked on the tail with non-toxic dye on the previous days. Then the tests were performed in the following order: social recognition test, urine marking test (males), single-caging, resident-intruder test (males). Video records were used to generate tracks representing the position of mice. Tracks were analyzed with the EthoVision software (Noldus Information Technology, Solingen, Germany)

#### Test for social novelty preference

The social paradigm was employed to examine general sociability and cognition in the form of social novelty preference. The test was performed as described (Muhia *et al*, 2022). A discrimination index used to assess social memory was calculated according to the following formula: Discrimination index = [(TimeNovel – TimeFamiliar)/TimeNovel+Familiar]. The calculation leads to a score ranging from −1 (preference for the familiar conspecific) to +1 (preference for a novel mouse). One-sample t-tests were used to compare the discrimination measure of each genotype group with zero (random exploration value). An additional analysis was conducted to compare the level of discrimination between genotype groups.

#### Urine marking test

The urine marking test was performed as previously described (Fellini & Morellini, 2013). In brief, three age-matched WT female mice were placed under a fenestrated plexiglass wall in a corner of the 50x50 cm arena. The focal male mouse was placed in the arena free to explore for 30 minutes. Urine markings were collected on grade 4 Whatman filter paper placed on the arena floor during the experiments and counted under UV light for each quadrant of the arena.

#### Resident intruder test

The resident intruder test was conducted according to a previous publication (Schob *et al*, 2019). Male focal mice were single-caged for 72 hours. The home cage was gently transferred from the vivarium to the experimental room. A plexiglas panel provided with ventilation holes substituted the cage top and the nesting material was completely removed. After 5 min, a C57Bl/6J age- and body-weight-matched unfamiliar male (intruder) was introduced into the cage of the focal animal (resident). The test lasted for 10 min after the first resident–intruder contact. Social interactions were analyzed using the Observer software blind to genotype. The experimenter trained himself until repeatedly scoring at least 80% consistency between two analyses of the same mouse performed at different times (calculated using the reliability test of the Observer software; maximal time discrepancy between two evaluations: 1 s).

## Acknowledgements

We thank Fabio Morellini for advice and support in data analysis. Supported by a German Research Foundation (DFG) grant KN556 to MK.

## References

Andreu Z, Yanez-Mo M (2014) Tetraspanins in extracellular vesicle formation and function. Front Immunol 5: 442

Bassani S, Cingolani LA, Valnegri P, Folci A, Zapata J, Gianfelice A, Sala C, Goda Y, Passafaro M (2012) The X-linked intellectual disability protein TSPAN7 regulates excitatory synapse development and AMPAR trafficking. Neuron 73: 1143–1158

Bassani S, Passafaro M (2012) TSPAN7: A new player in excitatory synapse maturation and function. Bioarchitecture 2: 95–97

Becic A, Leifeld J, Shaukat J, Hollmann M (2021) Tetraspanins as Potential Modulators of Glutamatergic Synaptic Function. Front Mol Neurosci 14: 801882

Charrin S, le Naour F, Silvie O, Milhiet PE, Boucheix C, Rubinstein E (2009) Lateral organization of membrane proteins: tetraspanins spin their web. Biochem J 420: 133–154

Drickamer LC (2001) Urine marking and social dominance in male house mice (Mus musculus domesticus). Behav Processes 53: 113–120

Fellini L, Morellini F (2013) Mice create what-where-when hippocampus-dependent memories of unique experiences. J Neurosci 33: 1038–1043

Freitag S, Schachner M, Morellini F (2003) Behavioral alterations in mice deficient for the extracellular matrix glycoprotein tenascin-R. Behav Brain Res 145: 189–207

Leroy F, Park J, Asok A, Brann DH, Meira T, Boyle LM, Buss EW, Kandel ER, Siegelbaum SA (2018) A circuit from hippocampal CA2 to lateral septum disinhibits social aggression. Nature 564: 213–218

Lozano R, Rosero CA, Hagerman RJ (2014) Fragile X spectrum disorders. Intractable Rare Dis Res 3: 134–146

Muhia M, YuanXiang P, Sedlacik J, Schwarz JR, Heisler FF, Gromova KV, Thies E, Breiden P, Pechmann Y, Kreutz MR et al (2022) Muskelin regulates actin-dependent synaptic changes and intrinsic brain activity relevant to behavioral and cognitive processes. Commun Biol 5: 589

Murru L, Moretto E, Martano G, Passafaro M (2018) Tetraspanins shape the synapse. Mol Cell Neurosci 91: 76–81

Murru L, Ponzoni L, Longatti A, Mazzoleni S, Giansante G, Bassani S, Sala M, Passafaro M (2021) Lateral habenula dysfunctions in Tm4sf2(-/y) mice model for neurodevelopmental disorder. Neurobiol Dis 148: 105189

Prox J, Willenbrock M, Weber S, Lehmann T, Schmidt-Arras D, Schwanbeck R, Saftig P, Schwake M (2012) Tetraspanin15 regulates cellular trafficking and activity of the ectodomain sheddase ADAM10. Cell Mol Life Sci 69: 2919–2932

Schob C, Morellini F, Ohana O, Bakota L, Hrynchak MV, Brandt R, Brockmann MD, Cichon N, Hartung H, Hanganu-Opatz IL et al (2019) Cognitive impairment and autistic-like behaviour in SAPAP4-deficient mice. Transl Psychiatry 9: 7

Seipold L, Altmeppen H, Koudelka T, Tholey A, Kasparek P, Sedlacek R, Schweizer M, Bar J, Mikhaylova M, Glatzel M et al (2018) In vivo regulation of the A disintegrin and metalloproteinase 10 (ADAM10) by the tetraspanin 15. Cell Mol Life Sci 75: 3251–3267

Stajano D, Lombino FL, Schweizer M, Glatzel M, Saftig P, Gromova KV, Kneussel M (2023) Tetraspanin 15 depletion impairs extracellular vesicle docking at target neurons. J Extracell Biol 2:e113

Takahashi A, Miczek KA (2014) Neurogenetics of aggressive behavior: studies in rodents. Curr Top Behav Neurosci 17: 3–44

van Deventer SJ, Dunlock VE, van Spriel AB (2017) Molecular interactions shaping the tetraspanin web. Biochem Soc Trans 45: 741–750

